## Supplementary information for "Characterization of a novel temperate phage facilitating *in vitro* dispersal of multicellular bacteria"

**Running title:** Phage-mediated dispersal of multicellular bacteria

### LIST:

**Supplementary Fig. S1**: Schematic representation of deletions generated by a CRIPSR-Cas9 approach targeting Samy prophage integrase

**Supplementary Fig. S2**: GC content (A) and size (B) of a panel of 330 *Streptomyces* phages

**Supplementary Fig. S3**: Multidimensional analyses of the transcriptomes of *S. ambofaciens* ATCC 23877 grown under various conditions

**Supplementary Fig. S4**: Heatmaps of pSAM1 (A) and pSAM2 (B) transcriptomes in different growth conditions

**Supplementary Fig. S5:** Samy expression profile under the different conditions studied

**Supplementary Fig. S6:** Samy phage morphology and infection assays

**Supplementary Fig. S7:** Results of high-throughput sequencing of the double-stranded DNA virome of *Streptomyces ambofaciens* ATCC 23877 grown 4 days in BM medium

**Supplementary Fig. S8:** Morphology of *S. ambofaciens* and *S. coelicolor* after 4 days growth in BM medium

**Supplementary Fig. S9:** Counting of colony forming units after 4 days of growth in BM medium of *S. ambofaciens* ATCC 23877 reference strain and its isogenic ΔSamy #clone 3 mutant supplemented with conditioned supernatants

**Supplementary Table S1**: Strains used in this study

**Supplementary Table S2**: Primers used in this study

**Supplementary Table S3**: Media used in this study

**Supplementary Table S4**: List and characteristics of 330 *Streptomyces* phages referenced in Actinophage Database, NCBI and/or ICTV. The legend is detailed in the “Readme” sheet.

**Supplementary Table S5**: Results from the OSMAC-RNAseq approach conducted in *Streptomyces ambofaciens* ATCC 23877. The legend is detailed in the “Readme” sheet.

**Supplementary Table S6**: List of Samy genes overexpressed in non or poorly inducing conditions. The legend is detailed in the “Readme” sheet.

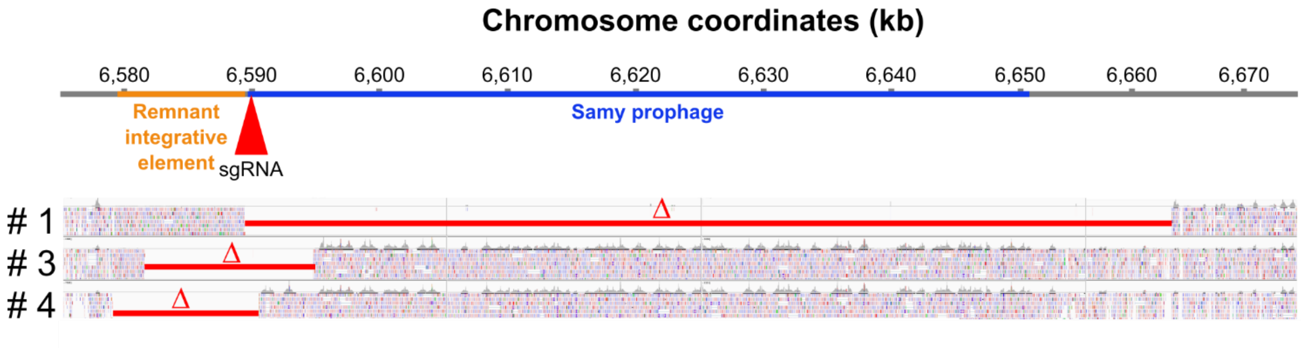


**Supplementary Fig. S1: Schematic representation of deletions generated by a CRIPSR-Cas9 approach targeting Samy prophage integrase**

The Samy integrase gene position targeted the sgRNA used in this study is indicated by a red triangle. The location of the remnant integrative element and Samy prophage are colored in orange and blue, respectively. The position of the deletion (Δ) observed in each clone is highlighted by red lines. The visualization of the position of sequencing reads obtained for each clone was generated using the Integrative Genome Viewer (IGV). The precise position of each deletion is detailed in the **Table S1**.

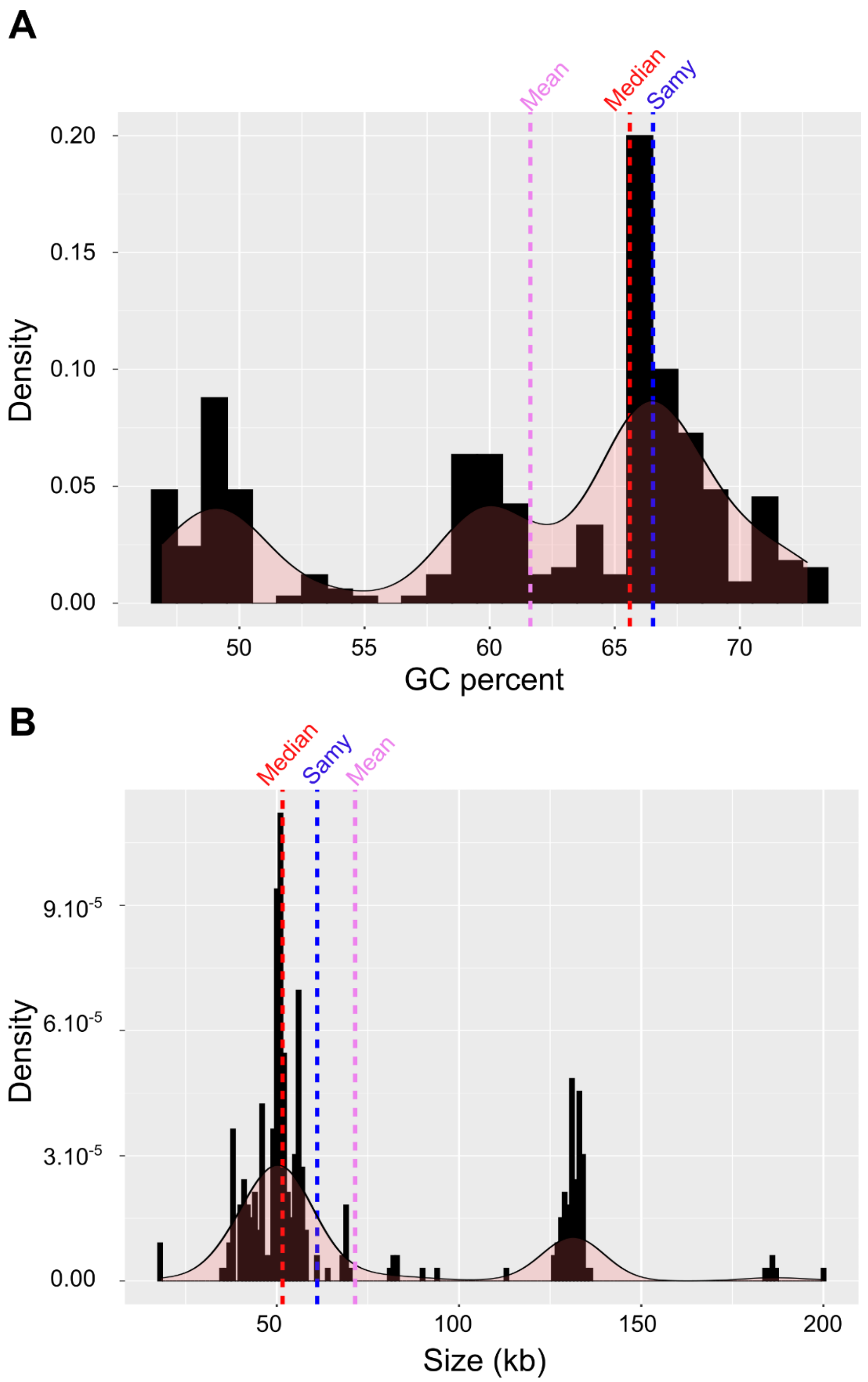


**Figure S2: GC content (A) and size (B) of a panel of 330 *Streptomyces* phages**

This analysis was conducted on a panel of 330 sequenced phages, including Samy, listed in the Actinophage Database, NCBI and/or ICTV databases (Supplementary **Table S4**). Since the distribution of values is multimodal, information about both the mean (pink dashed line) and median (red dashed line) was taken into account to compare to Samy (blue dashed line). The density plots were generated using a bin width of 1 and 1000 in panels A and B, respectively. The density curves are indicated in each panel.

**A**

| Condition name | Medium type | Medium composition | Time point (h) | GEO accession number |
|---|---|---|---|---|
| MP24 | Liquid | MP5 | 24 | GSE162865 |
| MP30 | Liquid | MP5 | 30 | GSE162865 |
| MP36 | Liquid | MP5 | 36 | GSE162865 |
| MP48 | Liquid | MP5 | 48 | GSE162865 |
| MP72 | Liquid | MP5 | 72 | GSE162865 |
| Y24 | Liquid | YEME 10% saccharose | 24 | GSE162865 |
| Ycongo24 | Liquid | YEME 10% saccharose + 1µg/ml congocidine | 24 | GSE162865 |
| Y48 | Liquid | YEME 10% saccharose | 48 | GSE162865 |
| MM | Solid | Minimal Medium 0.5 % Mannitol | 30 | GSE232795 |
| NAG | Solid | MM + 25 mM N-Acetyl-Glucosamine | 30 | GSE232795 |
| SAF | Solid | SAF | 30 | GSE232795 |
| ONA | Solid | Oxoid Nutrient Agar | 30 | GSE232795 |
| HT | Solid | Hickey-Tresner agar | 30 | GSE232795 |

**B**

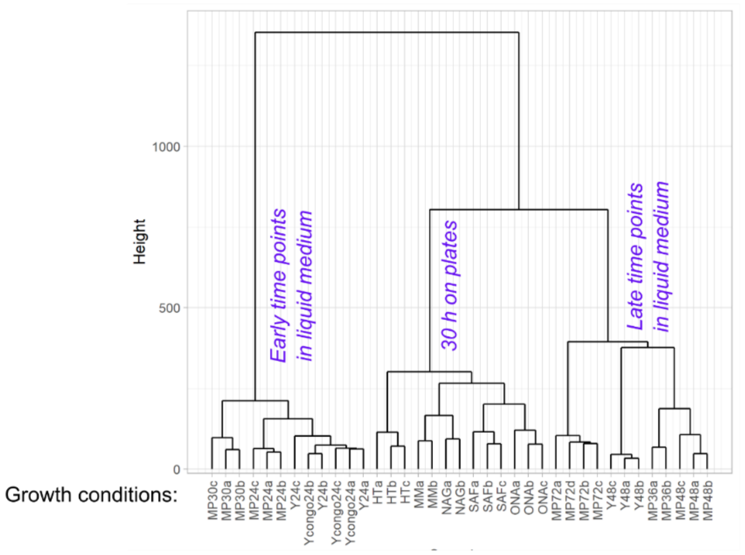


**C**

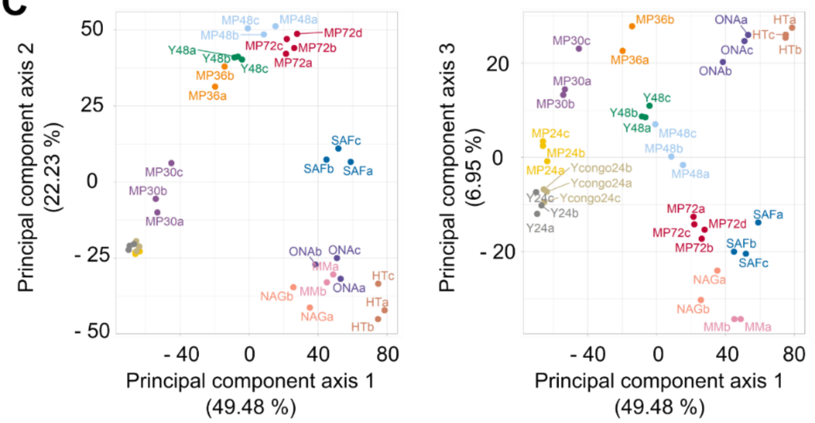


**Figure S3: Multidimensional analyses of the transcriptomes of *S. ambofaciens* ATCC 23877 grown under various conditions**

A. **Main characteristics of the various growth conditions analyzed in this study**.
B. **Ascending hierarchical classification** (Euclidean distance, Ward criterion) of the whole data set (37 samples). Transcriptomes form clusters based on biological conditions specified in blue near each node. The acronyms and colors indicate the condition (see panel A) with a distinct letter (a, b, c or d) for each replicate.
C. **Principal component analysis** (PCA) of the whole data set, with percentages of variance associated with each axis. The acronyms and colors indicate the condition (see panel A) with a distinct letter (a, b, c or d) for each replicate.

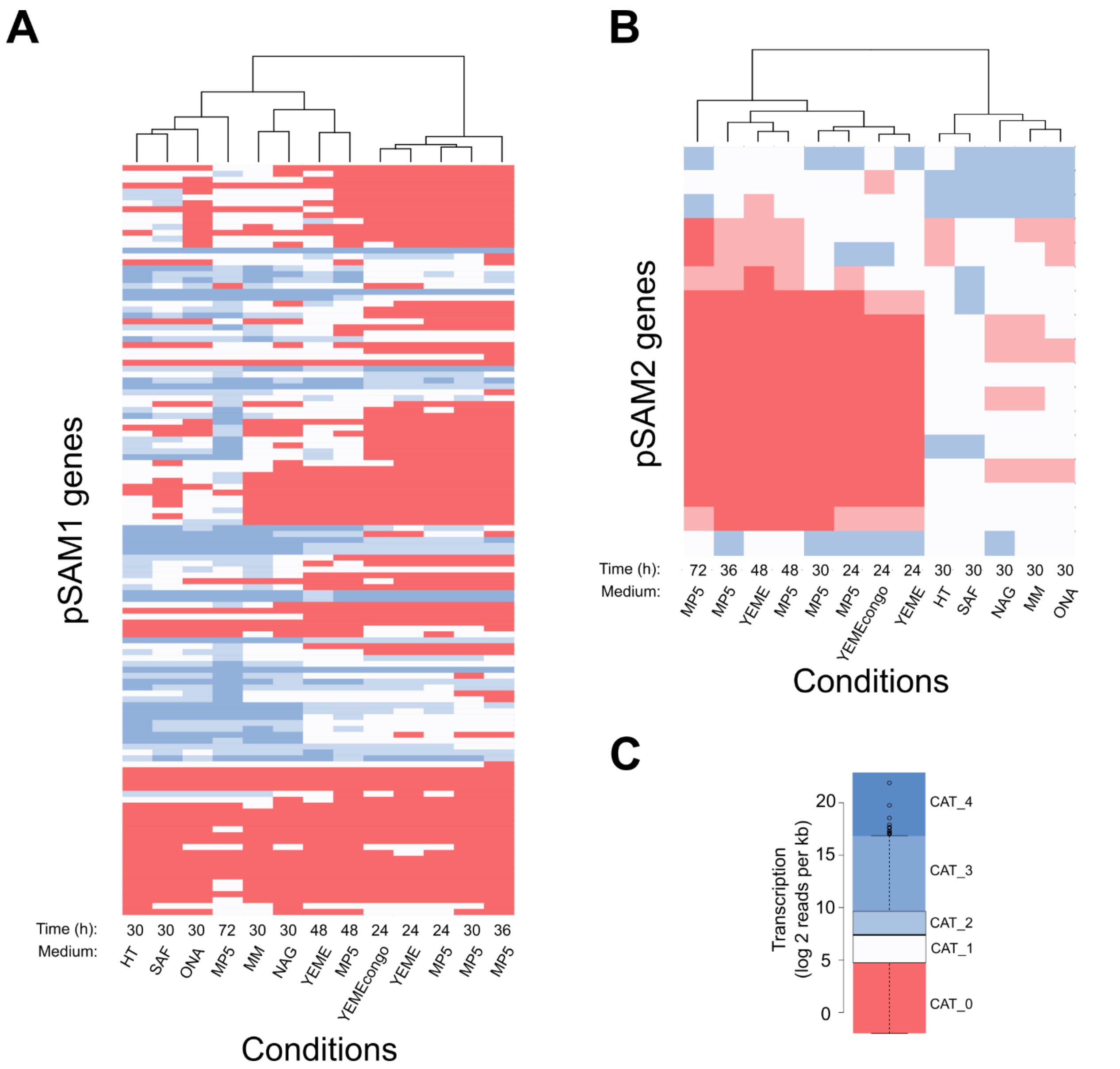


**Figure S4: Heatmaps of pSAM1 (A) and pSAM2 (B) transcriptomes in different growth conditions**

RNA-seq data (DESeq2 normalized number of reads per kb per gene, log2) were categorized in a color scale reflecting their expression relative to whole genome expression in each condition, as illustrated in panel C. Each line represents a gene, ranked according to order on the genome. Growth conditions (columns) are ranked by hierarchical clustering according to pSAM1 and pSAM2 transcriptomes, in panels A and B, respectively.

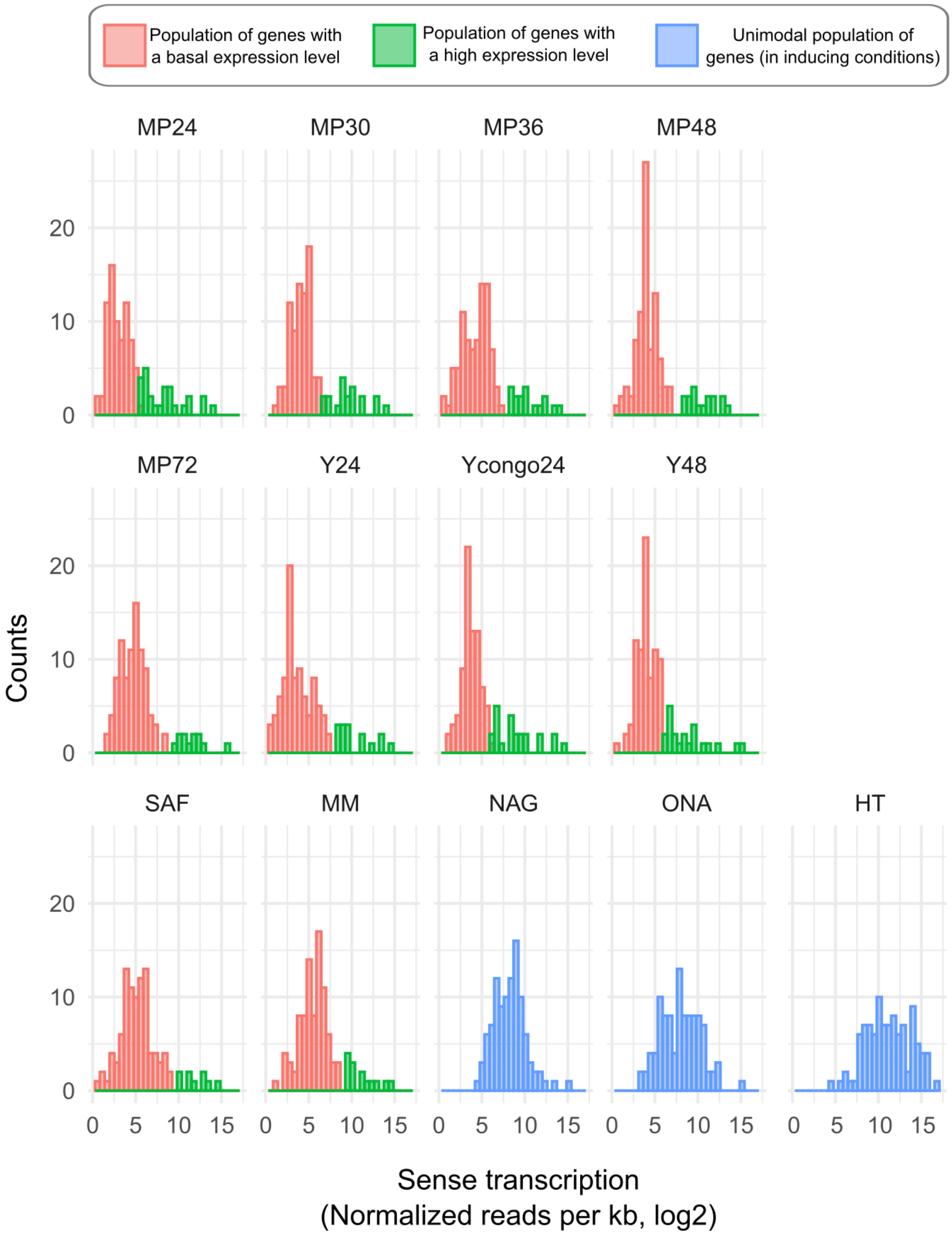


**Figure S5: Samy expression profile under the different conditions studied**

The expression level distribution of Samy genes (*n* = 102) is presented in each condition (described in Supplementary **Fig. S3** and **Table S3**). A dedicated R package ('mClust') was used to identify the bimodal distributions and classify genes accordingly. In non-inducing conditions (*i.e.* all conditions except HT, ONA, NAG), a bimodal profile was observed, genes being categorized in the basal (red) or high expression level (green) population. In 'HT', 'ONA' and 'NAG' conditions, the distribution of Samy gene expression was unimodal (population in blue).

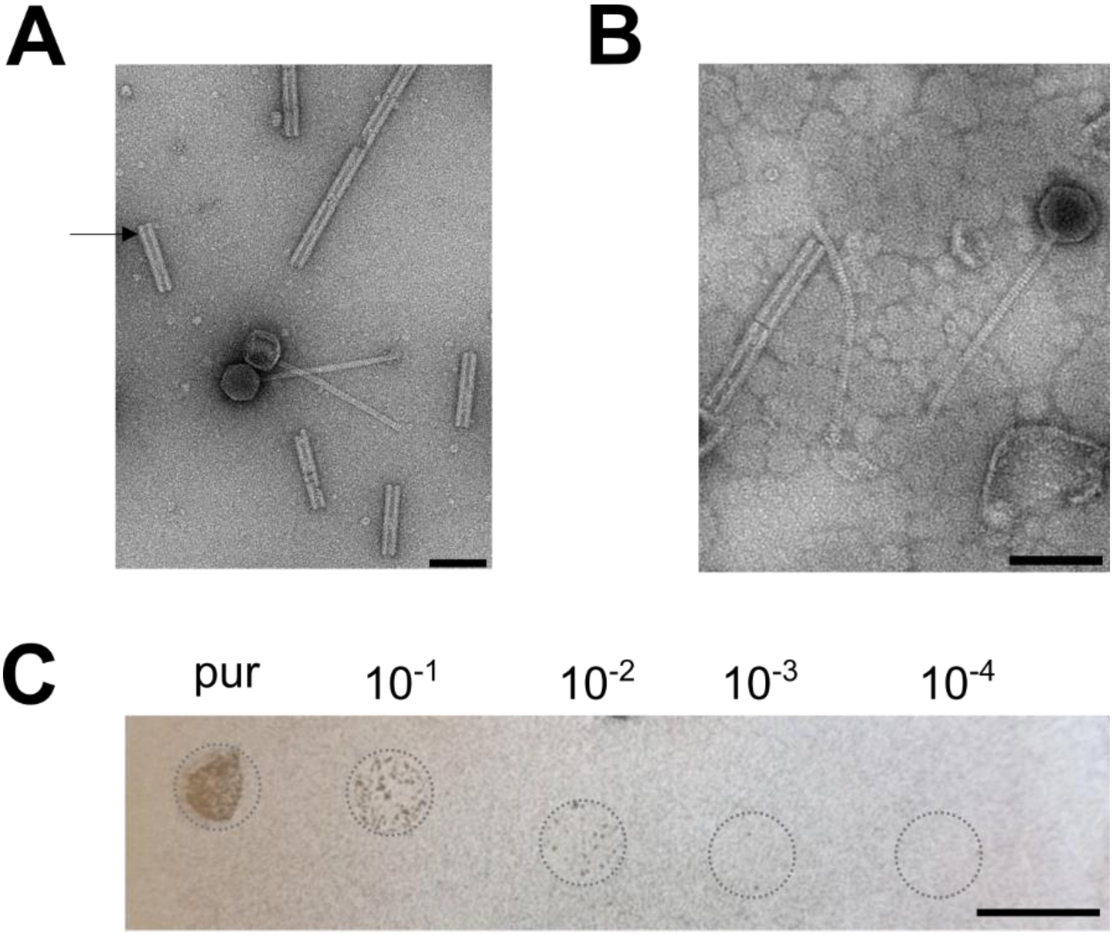


**Figure S6: Samy phage morphology and infection assays**

**A. Imaging of Samy phage produced in BM medium by transmission electron microscopy.** *S. ambofaciens* ATCC 23877 was grown during 4 days in BM medium. The supernatant was concentrated by CsCl-gradient ultracentrifugation. Viral particles were negatively stained with uranyl acetate. Two Samy phages surrounded by regular double tubular structures (arrow) appearing to originate from a larger structure, composed of subunits of these. These structure have been described as phage tail-like nanostructures and identified as extracellular contractile injection systems (1). Scale bar: 100 nm.

**B. Imaging of Samy phage produced in HT medium by transmission electron microscopy.** *S. ambofaciens* ATCC 23877 was grown during 3 days in HT medium. Supernatant was harvested and concentrated by an iodixanol gradient. The sample was negatively stained with uranyl acetate. Samy virions and extracellular contractile injection systems were also observed. Scale bar: 100 nm.

**C. Infection of *S. lividans* TK24 by Samy phage.** Five µl of serial dilutions of Samy phage produced after 4 days-growth in BM medium were spotted on a lawn of *S. lividans* TK24 spores poured onto an SFM plate. The picture was taken after one week of growth at 30°C. Spotted sites are framed by a grey circle. Viral plaques appear as small holes in the *S. lividans* lawn (white). Scale bar: 1 cm.

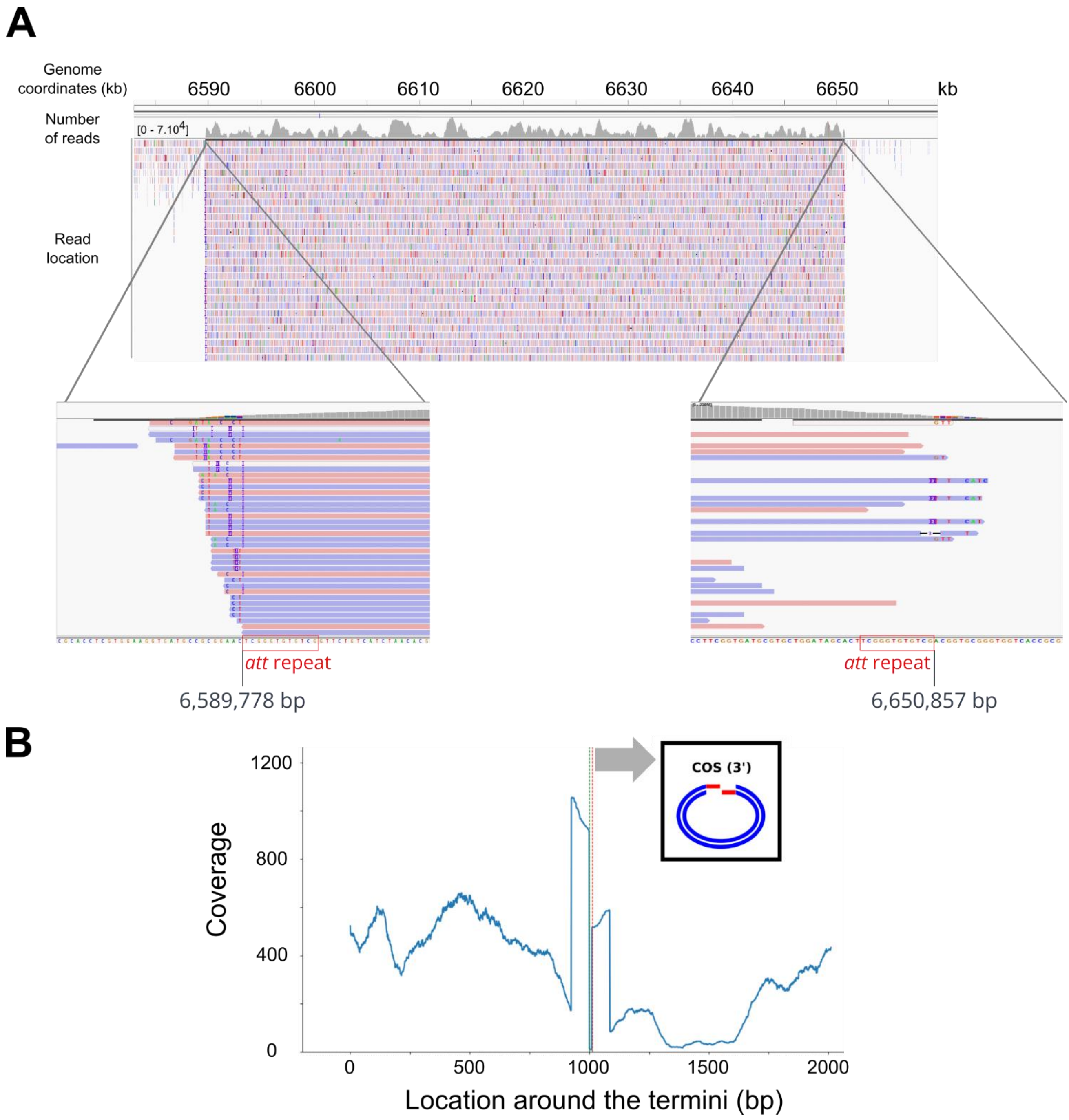


**Figure S7: Results of high-throughput sequencing of the double-stranded DNA virome of *Streptomyces ambofaciens* ATCC 23877 grown 4 days in BM medium**

**A. Sequence coverage around Samy prophage region.** The coordinates of *S. ambofaciens* ATCC 23877 chromosome are indicated. The inserts present the focus on the regions surrounding the repeats contained in the *att*L and *att*R sites. The data were visualized using the Integrative Genome Viewer (v2.8.0) software.

**B. Sequence coverage at termini positions identified by PhageTerm** (2). Exact termini positions are represented by dotted lines (Red: left; Green: right). The predicted cohesive sequence (*cos*) is: CGTTAAGGTGC (from 6,593,965 to 6,593,975 bp position on *S. ambofaciens* ATCC 23877 chromosome). The analysis was obtained by PhageTerm (2) run from the Galaxy server (https://galaxy.pasteur.fr).

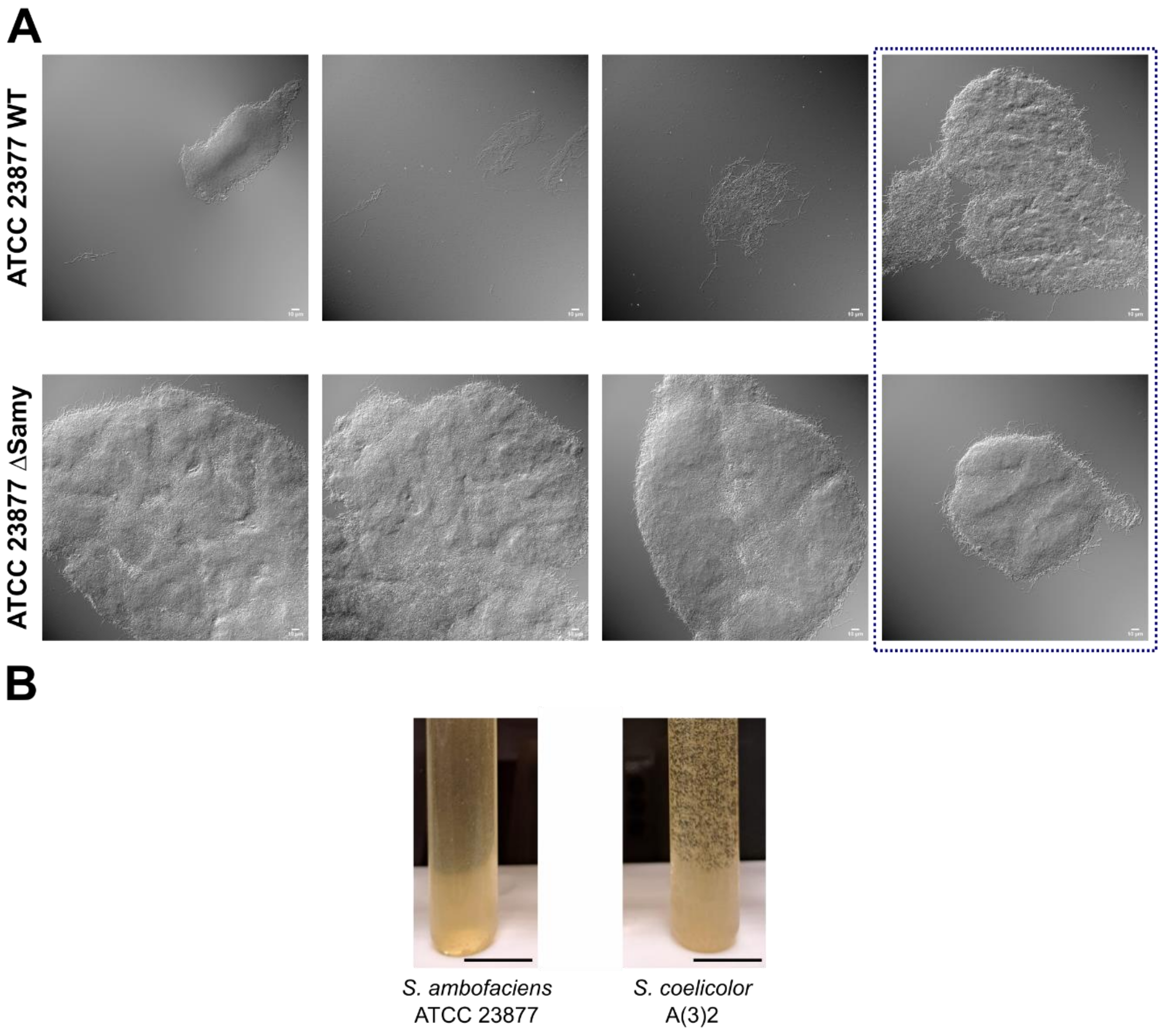


**Figure S8: Morphology of *S. ambofaciens* and *S. coelicolor* after 4 days growth in BM medium**

A. **Microscopy of *Streptomyces* aggregates**. *S. ambofaciens* ATCC 23877 and its derivate CRISPR-deleted of Samy prophage (clone #3) were grown in BM medium and imaged using differential interference contrast. The framed images present rare observations (clusters in the WT strain, small pellets in the Samy-deleted strain). Additional fields of the experiment presented in **Fig. 5.C** panel. Scale bar: 10 µm.
B. **Dispersed *versus* aggregated growth of *S. ambofaciens* ATCC 23877 and *S. coelicolor* A(3)2 strains in BM media.** The results are representative of the appearance of the most frequently observed cultures, as the quantity and size of cell clusters may vary from one experiment to the other. Scale bar: 1.5 cm.

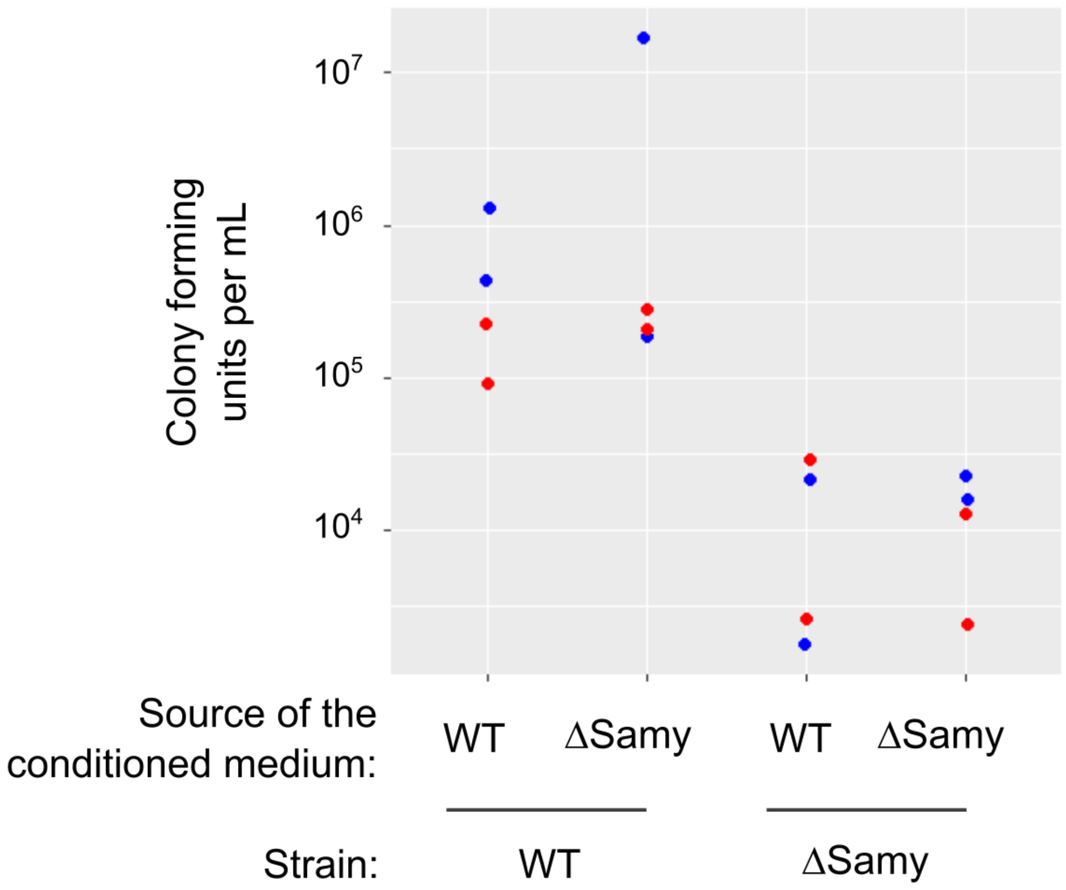


**Figure S9: Colony forming units after 4 days of growth in BM medium of *S. ambofaciens* ATCC 23877 reference strain and its isogenic ΔSamy clone #3 mutant supplemented with conditioned supernatants**

*S. ambofaciens* ATCC 23877 WT strain or its isogenic mutant ΔSamy clone #3 (Supplementary **Table S1**) were grown for 24 h before supplementing the medium with filtered conditioned supernatant from a previous 4 day culture of the WT or Samy-deleted strain at ratios of 1:2 (blue) or 1:5 (red) of conditioned supernatant per 24h-culture volumes. Then, bacteria were grown for an additional 3 days before counting. The results of 2 independent experiments are presented.

**Table S1: Strains used in this study***

| Name | Main characteristics | Samy complete prophage | Reference |
|---|---|---|---|
| Strains used to perform Samy infection assays | | | |
| *S. ambofaciens* ATCC 23877 | RP3486 strain deposited at the ATCC by RP - Sequenced genome (GCF_001267885.1_ASM126788v1) | + | (3), genome sequence (4), Pernodet-Lautru's lab collection |
| *S. ambofaciens* DSM 40697 | Tü13 strain deposited at the DSM collection - Sequenced genome (GCF_001632865.1, ASM163286v1) | - | (5), genome sequence (6), Pernodet-Lautru's lab collection |
| *S. albidoflavus* J1074/R2 | Derivate from *S. albidoflavus* J1074 | - | (8), Pernodet-Lautru's lab collection |
| *S. coelicolor* A(3)2 | Type strain | - | (9), Pernodet-Lautru's lab collection |
| *S. lividans 66* TK24 | Streptomycin-resistant mutation (*str6)* genetic marker | - | (10), Pernodet-Lautru's lab collection |
| *S. noursei* ATCC 11455 | *S. noursei* Brown *et al.* | - | (11), Pernodet-Lautru's lab collection |
| *S. rimosus* ATCC 10970 | Isolate from soil | - | (12), Pernodet-Lautru's lab collection |
| *S. venezuelae* ATCC 10712 | *S. venezuelae* Ehrlich *et al.,* type strain isolated from soil in Venezuela | - | Pernodet-Lautru's lab collection |
| Other *Streptomyces ambofaciens* parental strains | | | |
| *S. ambofaciens* RP3486 | Isolated from soil in France (Péronne, Somme) by RP | + | (3), Pernodet-Lautru's lab collection, initially obtained from RP |
| *S. ambofaciens* ETH 6703 | Isolated from soil in Italy (Rome) by H. Zähner, collection from the ETH | - | (5), Pernodet-Lautru's lab collection |
| *S. ambofaciens* Tü13 | Collection isolate of ETH 6703 from Tübingen university | - | (5), Pernodet-Lautru's lab collection |
| *S. ambofaciens* ETH 11317 | Collection from the ETH, initially named *S. aureofaciens* | - | (5), (7), Pernodet-Lautru's lab collection |
| Derivates of *Streptomyces ambofaciens* RP3486 | | | |
| *S. ambofaciens* NRRL 2420 | *S. ambofaciens* Pinnert-Sindico (ATCC 15154) also named 'Rhone-Poulenc 1297-18-T2, | + | Pernodet-Lautru's lab collection |

| | | | |
|---|---|---|---|
| | obtained by mutagenesis (UV irradiation) of RP3486 | | |
| *S. ambofaciens* JI3212 | Derivate of ATCC 15154 maintained by the John Innes Centre (the strain harbors a mutation in pSAM2 compared to ATCC 15154) | + | (13), Pernodet-Lautru's lab collection, initially obtained from the John Innes Institute collection |
| *S. ambofaciens* RP181110 | Isolated after UV irradiation of RP3486 | + | (13), Pernodet-Lautru's lab collection, initially obtained from RP |
| Derivates of *Streptomyces ambofaciens* ATCC 23877 | | | |
| *S. ambofaciens* ΔSamy clone #1 | Isolated after CRISPR-Cas9 engineering with a sgRNA targeting Samy integrase encoding gene. Harbors a large deletion (74.0 kb) compassing Samy and ≈ 12.7 kb downstream sequences (Δ6,589,492-6,663,570) | - | This study |
| *S. ambofaciens* ΔSamy clone #3 | Isolated after CRISPR -Cas9 engineering with a sgRNA targeting Samy integrase encoding gene. Harbors a local deletion (≈ 13.2 kb) of the region surrounding Samy integrase gene and the remnant integrative element located upstream Samy (Δ6,581,646-6,594,871) | - | This study |
| *S. ambofaciens* ΔSamy clone #4 | Isolated after CRISPR -Cas9 engineering with a sgRNA targeting Samy integrase encoding gene. Harbors a deletion (≈ 11.5 kb) of Samy integrase gene and all of the remnant integrative element located upstream Samy (Δ6,579,099-6,590,590) | - | This study |

*Abbreviations: ATCC (American Type Culture Collection), DSM (*Deutsche Sammlung von Mikroorganismen*), ETH (*Eidgenössische Technischen Hochschule*, Zürich), RP (Rhône-Poulenc)

**Table S2: Primers used in this study**

| Name | Sequence (5'-3') | Target |
|---|---|---|
| Quantitative PCR | | |
| SBM399 | CCGCTGACCTCGTCGTCTAC | Region 6,617,269-6,617,375 of *S. ambofaciens* ATCC 23877 chromosome corresponding to a Samy gene (SAMYPH_28/ SAM23877_RS29055) encoding a tail fiber protein |
| SBM400 | GCCCTTGATGGTGTTCAGGA | |
| SBM446 | CGAAGCCGCTCAGGCCAACC | Region 5,975,814-5,975,937 of *S. ambofaciens* ATCC 23877 chromosome containing part of the *srmS* (*srm40*, SAM23877_RS26595) *S. ambofaciens* ATCC 23877 gene encoding the regulator of the spiramycin BGC – This qPCR was used as a control to test the presence of host DNA after DNAse treatment. |
| SBM447 | GCCCACCCGCACCATGAAGG | |
| Primers used to clone the sgRNA targeting Samy integrase | | |
| SBM393-sgRNA – int2 | CATG**CCATGG**ACACATCGACGATGGTAGGTGTTTTAGAGCTAGAAATAGC | *Nco*I-*Sna*BI sgRNA cloning site of pCRISPR-Cas9 vector from Tong *et al*.(14) |
| SBM67-scaffold-R | ACGCC**TACGTA**AAAAAAGCACCGACTCGGTGCC | |

### Table S3: Media used in this study

| Name | Composition | Reference |
|---|---|---|
| Classical *Streptomyces* media | | |
| HT (Hickey and Tresner) | 1 g/L yeast extract, 1 g/L meat/beef extract (Difco™ Beef extract, ref 212610), 2 g/L bactotryptone, 10 g/L white dextrin, 0.02 g/L $CoCl_2$, with or without 20 g/L agar; pH 7.3 | Adapted from (15), (16) |
| MM (Minimal Medium Mannitol) | 0.5 g/L L-asparagine, 0.5 g/L $K_2HPO_4$, 0.2 g/L $MgSO_4$, 0.001 g/L $FeSO_4$, 0.5 % mannitol, 10 g/L agar; pH 7.0/7.2 | (15) |
| MP5 (medium of production n°5) | 7 g/L yeast extract, 20.9 g/L MOPS, 5 g/L NaCl, 1 g/L $NaNO_3$, 36 mL/l glycerol; pH 7.5 | (17) |
| ONA (Oxoid nutrient agar) | 28g / L of Oxoid nutrient agar (corresponding to 1 g/L of meat/beef extract, 2g/L yeast extract, 5g/L pepton, 5 g/L NaCl, 15 g/L agar, pH7.4) | CM003 (Oxoid) |
| SAF | 0.5 g/L yeast extract, 0.5 g/L meat/beef extract, 21 g/L MOPS, 5g/L glucose, 1 g/L enzymatic hydrolysate of casein, 20 g/L agar; pH 7.0 | (18) |
| SNA (Soft Nutrient Agar) | 2 g/L agar, 8 g/L nutrient broth MPBiomedicals (Cat#1007917) | (15) |
| SFM (Soy Flour-Mannitol) | 20g/L organic soy flour, 20 g/L mannitol, 20 g/L agar | (15) |
| Variants of the HT or MP5 media to generate the results presented in the Fig. 3 | | |
| HT - dextrin | 1 g/L yeast extract, 1 g/L meat/beef extract, 2 g/L bactotryptone, 0.02 g/L $CoCl_2$; pH 7.3 | This study |
| HT - meat | 1 g/L yeast extract, 2 g/L bactotryptone, 10 g/L white dextrin, 0.02 g/L $CoCl_2$; pH 7.3 | This study |
| HT – $CoCl_2$ | 1 g/L yeast extract, 1 g/L meat/beef extract, 2 g/L bactotryptone, 10 g/L white dextrin; pH 7.3 | This study |
| HT - dextrin – meat - $CoCl_2$ | 1 g/L yeast extract, 2 g/L bactotryptone; pH 7.3 | This study |
| HT - dextrin – meat | 1 g/L yeast extract, 2 g/L bactotryptone, 0.02 g/L $CoCl_2$; pH 7.3 | This study |
| BM (bacteriophage production medium) | 1 g/L yeast extract, 1 g/L meat/beef extract, 2 g/L bactotryptone; pH 7.3 | This study |
| BM + MOPS | 1 g/L yeast extract, 1 g/L meat/beef extract, 2 g/L bactotryptone, 21 g/L MOPS; pH 7.3 | This study |
| MP5 - MOPS | 7 g/L yeast extract, 5 g/L NaCl, 1 g/L $NaNO_3$, 36 ml/l glycerol; pH 7.5 | This study |
